# DivIVA phosphorylation at threonine affects its dynamics and cell cycle in *Deinococcus radiodurans*

**DOI:** 10.1101/2022.06.10.495630

**Authors:** Reema Chaudhary, Swathi Kota, Hari S. Misra

## Abstract

RqkA, a γ radiation responsive Ser/Thr quinoprotein kinase, is characterized for its role in radioresistance in *Deinoocccus radiodurans*. DivIVA is a cell division protein involved in determination of cell pole and division site in bacteria. RqkA phosphorylated cognate DivIVA (drDivIVA) at Threonine 19 (T19) residue located in its pole recognition motif. The phospho-mimetic replacement of T19 (T19E) functioned differently than phospho-ablative (T19A) and drDivIVA proteins. T19E-RFP expressing in wild type background showed arrest in dynamics of drDivIVA, and loss of interaction with genome segregation protein. *divIVA* is shown to be an essential gene in this bacterium. The allelic replacement of *divIVA* with T19E-RFP was not tolerated unless drDivIVA was expressed episomally while there was no effect of this replacement with T19A-RFP and drDivIVA-RFP. These results suggested that the T19 phosphorylation in drDivIVA by RqkA has affected in vivo functions of DivIVA that would render cell cycle arrest in this bacterium.

## Introduction

Proteins undergo several types of post translational modifications (PTMs) *albeit* majority of them have been reported in eukaryotes. Amongst these, the most common PTMs are phosphorylation, acetylation, methylation, carbonylation, glycosylation and ubiquitination (Witze et al., 2007). The phosphoprotein homeostasis depends upon the active presence of cognate kinases and phosphatases in the cell. Different types of protein phosphorylation have been reported in different organisms, and the selectivity in their functions has been shown in different organisms. For example, Ser/Thr/Tyr phosphorylation are predominately studied in the DNA damage response and cell cycle regulation in eukaryotes, the histidine kinases as the part of two component systems, are best characterized as stress response kinases in bacteria (Chang & Stewart, 1998; Galperin & Nikolskaya, 2001; Mascher, 2006). However, understanding the roles of Ser/Thr protein kinases (STPKs) in the maintenance of genome integrity and regulation of cell division has also started growing in bacteria. For instances, the eukaryotic type STPKs was first identified in the Gram-negative bacterium *Myxococcus xanthus* (Munoz-Dorado et al., 1991). Recently, STPKs have been characterized from both biotic and abiotic stress tolerant bacteria including pathogens like *Pseudomonas aeruginosa* (Mougous et al., 2007), *Enterococcus faecalis* (Hall et al., 2013), *Staphylococcus aureus* (Burnside et al., 2010), *Mycobacterium tuberculosis* (Av-Gay and Everett, 2000), *Streptococcus pneumoniae*, (Echenique et al., 2004), *Streptococcus agalactiae* (Rajagopal et al., 2003), *Streptococcus pyogenes* (Agarwal et al., 2011), and in super bug like *Deinococcus radiodurans* (Pereira et al., 2011; Rajpurohit et al, 2022). eSTPKs are promiscuous in nature that makes one STPK to phosphorylate several proteins. The Ser/Thr (S/T) phosphorylation has been reported in proteins involved in several biological processes including genome functions, virulence and stress response (Liu et al., 2011; Mata-Cabana et al., 2012; Ruggiero et al., 2012, Rajpurohit et al., 2022). The S/T phosphorylation has also been detected in bacterial cell division proteins and the effects of such phosphorylation on the functions of FtsZ have been characterized in many bacteria (Manuse et al., 2016, Grangeasse, 2016). DivIVA undergoes phosphorylation by StkP in *S. pneumoniae* and *S. suis* and the effects of such phosphorylation on the regulatory roles of this protein in cell division and morphogenesis have been reported (Giefing et al., 2008; Nováková et al., 2010, Ni et al, 2018). In *Mycobacterium*, Wag31 (a homolog of DivIVA) is involved in septum formation, cell wall synthesis and/or chromosome segregation and is found to be a substrate for protein kinases PknA and PknB (Witze et al., 2007).

*D. radiodurans* is characterized for its extraordinary resistance to DNA damaging agents including radiations and desiccation (Cox and Battista, 2005, Slade and Radman, 2011; Misra et al., 2013). The growth of this bacterium gets arrested upon γ radiation exposure and the levels of some cell division proteins do not change during post irradiation recovery (Modi et al., 2014). Interestingly, the RecA/LexA mediated canonical SOS response, which is the synonym to the DNA damage response and cell cycle regulation in majority of bacteria, is lacking in this bacterium (Little and Mount, 1982; Narumi et al., 2001, Bonacossa de Almeida et al., 2002). However, it modulates its proteome and transcriptome in response to DNA damage (Liu et al, 2003, Joshi et al., 2004). A few mechanisms that partly complement the absence of LexA/RecA mediated DNA damage response and cell cycle regulation have been suggested. For instance, DdrO/PprI (IrrE) regulating genome function by mimicking LexA/RecA model (Earl et al., 2002; Devigne et al., 2015; Narasimha and Basu, 2021), and guanine quadruplex (G4) DNA structure dynamics regulating genome function in response to γ radiation damage (Beaume et al., 2013; Kota et al., 2015, Mishra et al., 2019) have been suggested. These examples could explain the DNA damage responsive regulation of expression of those DNA repair proteins that have respective regulatory signatures. However, these mechanisms could not explain the regulation of function of cell division proteins during post irradiation recovery period of this bacterium. Independently, a DNA damage responsive Ser/Thr quinoprotein kinase (RqkA) has been characterized for its role in radiation resistance and double-strand break (DSB) repair in *D. radiodurans* (Rajpurohit and Misra, 2010). RqkA finds a number of putative substrates including PprA, RecA, DnaA, Hu, FtsZ, FtsA, MinC, FtsK and DivIVA in the proteome of this bacterium (Misra et al., 2013). The effect of S/T phosphorylation on the activity regulation of RecA, PprA and FtsZ have been demonstrated (Rajpurohit and Misra, 2013; Rajpurohit et al., 2016, Maurya et al., 2018). DivIVA of *D. radiodurans* (drDivIVA) has been functionally characterized for its interaction with cell division and genome maintenance proteins, and interestingly found to be an essential gene in this bacterium (Chaudhary et al., 2019; 2021). drDivIVA phosphorylation by RqkA and its essential nature made it an interesting protein to understand if its phosphorylation renders the cell to undergo growth arrest in response to DNA damage. Here, for the first time, we report the phosphorylation of drDivIVA by RqkA and demonstrated its significance in the regulation of drDivIVA functions by an allelic replacement of wild type with phospho-mimetic mutant alleles, and *in vivo* localization studies. We demonstrated that drDivIVA undergoes phosphorylation by a DNA damage responsive STPK (RqkA) and its subcellular role in the spatial regulation is arrested when phosphorylation at T19 was mimicked with glutamate, suggesting the possible involvement of drDivIVA phosphorylation in cell cycle arrest in response to γ radiation in *D. radiodurans*.

## Results

### RqkA phosphorylates its cognate drDivIVA at Threonine 19 (T19) residue

DivIVA is amongst other deinococcal proteins that possess the conserved motif “S/T-Q-X-hydrophobic-hydrophobic” (where X is any amino acid except the positively charged residues) predicted as the site of phosphorylation by eSTPK (as RqkA). The recombinant drDivIVA incubated with RqkA in the presence of [γ-^32^P] ATP produced 2 phosphobands, one each corresponding to drDivIVA and the auto phosphorylated form of RqkA (Fig 1A). drDivIVA phosphorylation by RqkA was further confirmed by *ex vivo* in transgenic *E. coli* BL21 cells co-expressing both RqkA and DivIVA (Fig 1B, C) and *in vivo* (Fig 1D). For *ex vivo*, the recombinant drDivIVA was purified from two backgrounds of *E. coli* BL21 cells, one expressed only DivIVA (pETD4A) and another co-expressed RqkA (pRadRqkA) and DivIVA (pETD4A). The phosphorylation status of the protein was checked using polyclonal anti-phospho Ser/Thr epitopes antibody (Cell Signaling, Inc). The recombinant RecA-P purified from the cells co-expressing RqkA was used as control. The DivIVA-P and RecA-P co-expressed in RqkA background showed phosphorylation while drDivIVA obtained from *E. coli* cells harboring only pETD4A did not show S/T phosphorylation (Fig 1C). Similarly, the phosphorylation of drDivIVA was checked in *D. radiodurans* cells expressing the tagged DivIVA episomally. The protein was checked for phosphorylation using antibodies against Ser/Thr epitopes and parallel confirmed for the protein identity using anti-drDivIVA antibody (Fig 1D). These results confirmed that DivIVA undergoes phosphorylation in *D. radiodurans*.

**Fig. 1:**
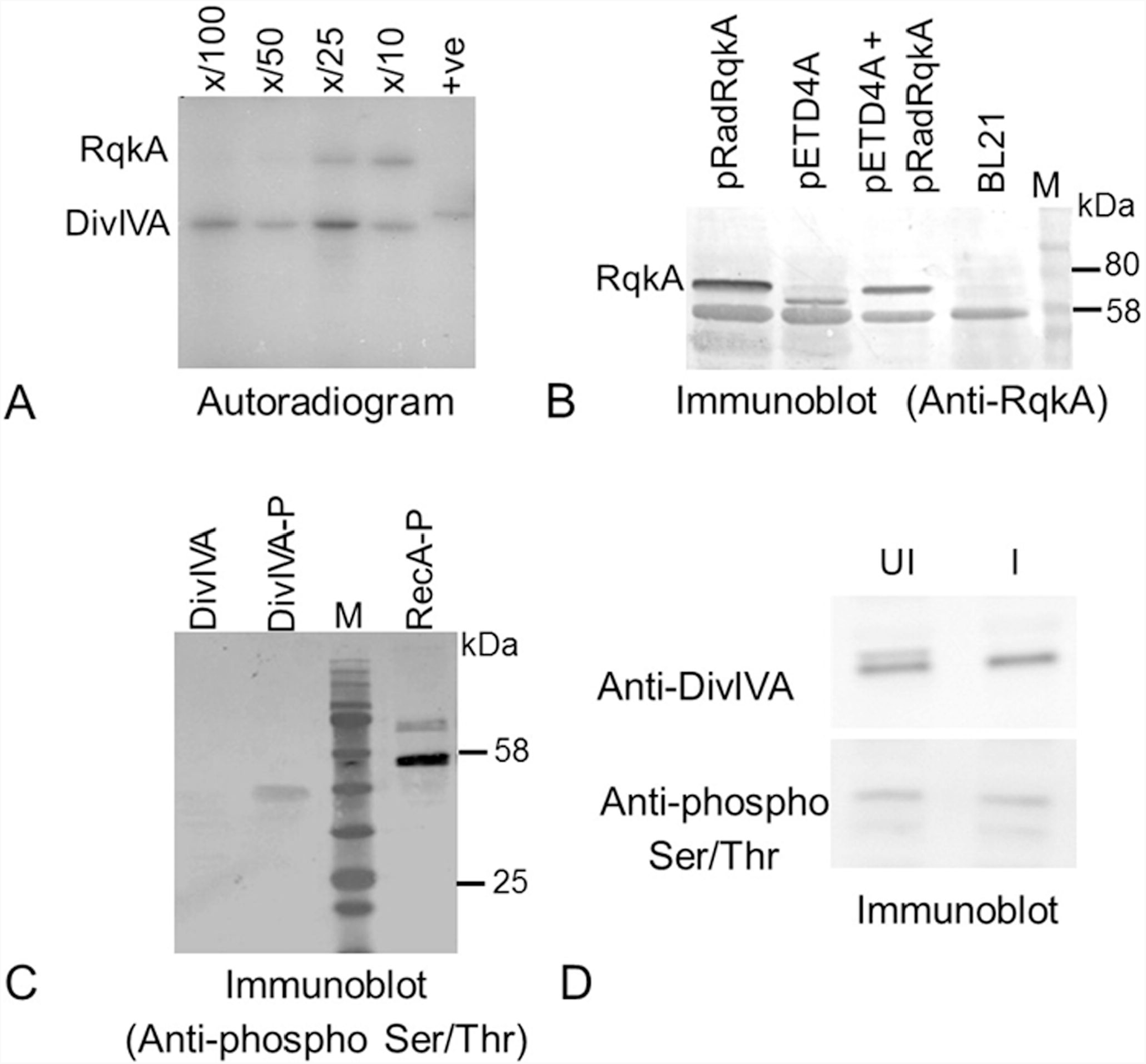
Phosphorylation of DivIVA by RqkA. DivIVA was incubated with increasing concentration of RqkA in kinase buffer containing [γ-^32^P] ATP and the reaction mixtures were separated on 12% SDS-PAGE and autoradiogram was developed **(A)**. *E. coli* BL21 cells expressing RqkA (pRadRqkA) and DivIVA (pETD4A) separately and together (pRadRqkA + pETD4A) in BL21 host (BL21). The expression of RqkA was confirmed by immunoblot using RqkA antibodies **(B)**. The DivIVA routed through RqkA expressing cells (DivIVA-P) and vector horboring cells (DivIVA) were run on SDS-PAGE and checked for phosphorylation signal using anti-phospho serine/threonine antibodies. The RecA routed through RqkA cells was used as +ve control (RecA-P) **(C)**. *In-vivo* phosphorylation of DivIVA was also checked in *D. radiodurans* harbouring pRGhisD4A. The deinococcal cells were grown and exposed to 6 kGy γ-radiation. The lysates of both unirradiated (UI) cells stored on ice and irradiated (I) cells were run on 12% SDS-PAGE and immunoblotted for both anti-DivIVA and anti-phospho Ser/Thr antibodies (D). Data shown is representative of a reproducible experiment repeated 3 times.

The phosphorylated form of drDivIVA was purified from *E. coli* BL21 cells co-expressing with RqkA and its non-phospho version from the cells harboring only expression vector (pETD4A). Both the proteins were subjected to mass spectrometric analysis (Fig S1A-B). As expected, no phosphoresidue(s) were detected with non-phospho preparation of drDivIVA. The mass spectrometric analysis detected a single phosphorylation site at Threonine 19 (T19) in drDivIVA routed through surrogate *E. coli* cells expressing RqkA *in trans*. Earlier, RecA and PprA showing very high levels of phosphorylation by RqkA were detected with multiple sites of phosphorylation (Rajpurohit et al, 2013 and 2016).

In *B. subtilis*, the residues Arginine-18 and Glycine-19 play a role in the targeting of DivIVA to the cell poles and hence constitute a polar-determining motif (Perry and Edwards, 2003). To understand the significance of T19 phosphorylation of drDivIVA, this phosphorylation motif was analyzed by the pair-wise alignment with DivIVA of *B. subtilis*. Interestingly, the N-terminal sequence of drDivIVA was found to be largely conserved with its numeral value of 9, and the T19 residue was a part of the polar determining motif of this protein (Fig S1C). Thus, a possibility of T19 having a role in pole determination in cocci and the effect of phosphorylation on its function could be hypothesized and investigated. Subsequently, the T19 residue in drDivIVA was replaced with alanine (hereafter referred as T19A) and glutamic acid (hereafter referred as T19E) and these phospho-mutants were checked for phosphorylation by RqkA *ex-vivo*. Results showed that both T19A and T19E mutants were phosphor-negative when incubated with RqkA (Fig S1E). These results provided strong evidence that drDivIVA is a Ser/Thr type phosphoprotein in *D. radiodurans* and is phosphorylated by RqkA at T19 residue.

### Wild type pattern of drDivIVA localization gets altered in T19E mutant in *D. radiodurans*

To understand the effect of T19 phosphorylation on the cellular dynamics of drDivIVA, *in vivo* localization of both wild type and phosphomutants of drDivIVA was monitored with respect to FtsZ. The C-terminal RFP fusion of drDivIVA (DivIVA-RFP) and its mutants, and GFP fusion of FtsZ (FtsZ-GFP) were co-expressed in *D. radiodurans* R1. The localization patterns of both the proteins were examined by confocal microscopy (Fig 2A-B). The pattern of FtsZ localization was nearly similar in all three cells types expressing drDivIVA or its phospho-mutants episomally in wild type background. For instances, the FtsZ ring was seen in closed conformation in nearly 48.27 ± 5.41 % to 50.1 ± 4.10 % cell population while 43.34 ± 4.89 % to 57.17 ± 5.72 % cell population had the FtsZ ring in an opened conformation in all three cell types (DivIVA^WT^, DivIVA^T19A^ and DivIVA^T19E^). It may be mentioned that all three cell types have wild type copy of DivIVA and if that has contributed to nearly no change in the localization of FtsZ pattern cannot be ruled out.

**Fig. 2:**
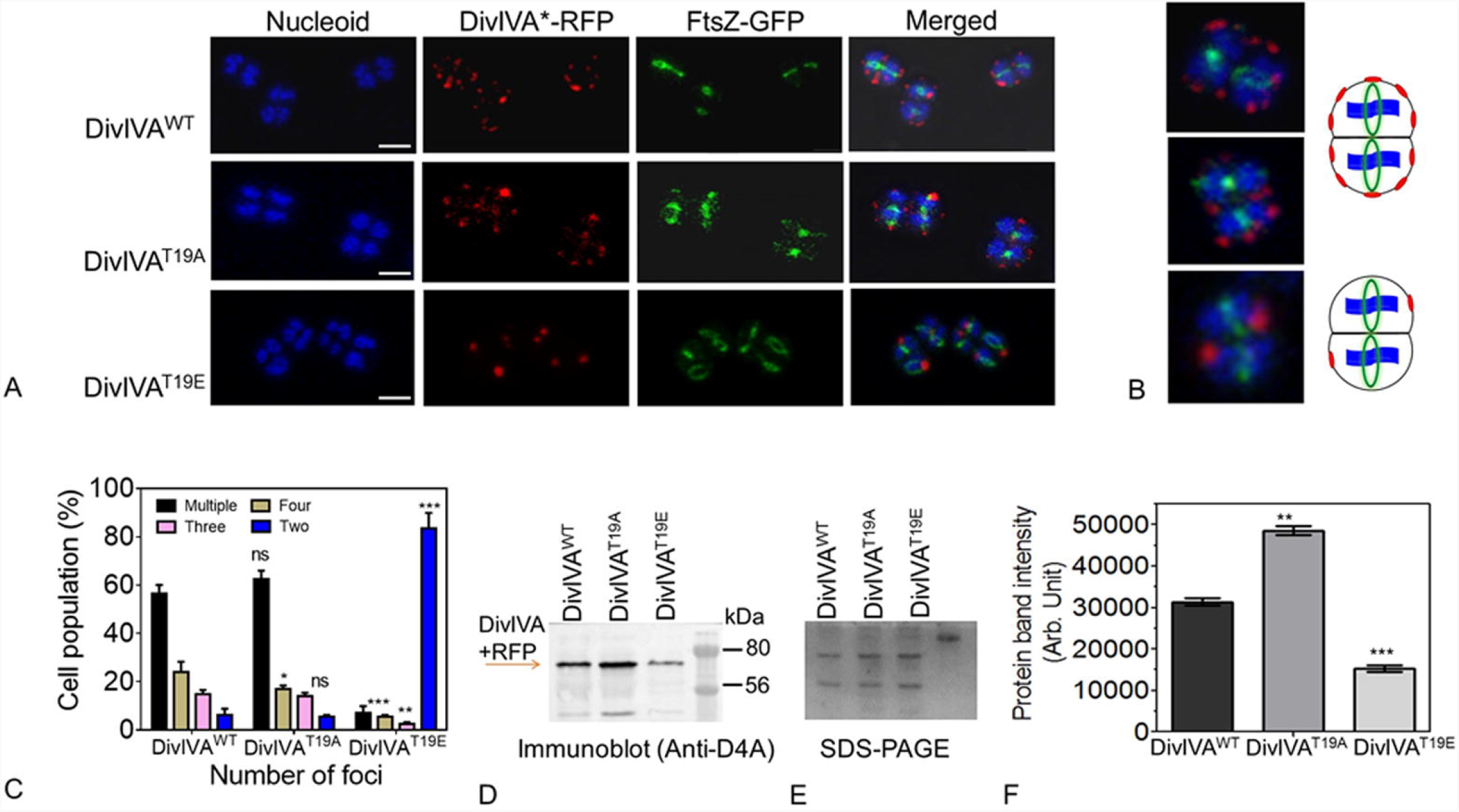
Localization of wild-type and phospho-mutant forms of DivIVA_Dr_ in *D. radiodurans*. Fluorescence imaging of *D. radiodurans* cells expressing DivIVA^WT^, DivIVA^T19A^ and DivIVA^T19E^ under constitutive promoter on pRadgro plasmid. The cells were observed for nucleoid staining (blue) and FtsZ-GFP localization (green) and DivIVA*-RFP where star indicates DivIVA variants (DivIVA^WT^, DivIVA^T19A^ and DivIVA^T19E^) and merged panel shows the image combining all the three channels (**A**). The zoom version of the representative cells are shown for clarity (**B**). These cells were counted for showing different types of foci (number of foci) **(C)**. The total proteins of *D. radiodurans* R1 harbouring pRGD4A (DivIVA^WT^), pRGD4A^T19A^ (DivIVA^T19A^) and pRGD4A^T19E^ (DivIVA^T19E^) were separated on SDS-PAGE (**E**) and immunoblotted with anti-DivIVA antibodies **(D)**. The intensity of protein bands was measured using Image J and plotted using Graphpad software **(F)**. Statistical analyses were done using Student *t*-test and the P values ≤0.01 and ≤0.005 are marked as (**) and (***), respectively. Data shown without statistical analysis is the representative of reproducible results repeated at least three times.

As far as the localization pattern of drDivIVA variants are concerned, the multiple DivIVA^WT^ foci were seen at the cell periphery and FtsZ foci were formed in the space unoccupied by DivIVA suggesting its role in the spatial positioning of FtsZ ring in *D. radiodurans*. Careful analysis showed that when the number of DivIVA foci was less than four, the cells would have committed for cell division. Nearly 56.5 ± 3.536% of the cells with multiple foci showed foci placements in juxtaposed fashion in the two compartments of the tetrad indicating that the cells are synchronized in the dynamics of this cell division protein. Nearly 62.5 ± 3.965% of cells expressing DivIVA^T19A^-RFP protein showed more than 4 foci and 14 ± 1.414% with less than four foci while majority (83.6 ± 6.364%) of cells expressing DivIVA^T19E^-RFP protein showed two foci per cell (Fig 2C). Although, the cumulative intensity of T19E foci appeared different than the cumulative intensity of T19A and drDivIVA foci separately, the possibility of this phenotype is due to the change in the levels of DivIVA variants was argued. The levels of DivIVA protein in the cells expressing drDivIVA and mutants (T19A and T19E) were measured by immunoblotting using protein specific antibodies (Fig 2D-E). Interestingly, the levels of episomally expressed drDivIVA and its mutant forms were found to be different. For instance, T19E protein was found to be less as compared to wild type and T19A. The difference in the levels of drDivIVA variants despite being under an identical promoter, translation machinery and translation fusion, is intriguing and a possibility of phosphorylation mediated homeostasis of drDivIVA in *D. radiodurans* could be speculated, and would be worth studying independently. Nevertheless, these results suggested that drDivIVA phosphorylation by RqkA at T19 site affects cellular localization of this protein in *D. radiodurans*.

### Mimicking T19 phosphorylation in drDivIVA affected its spatial placement with respect to FtsZ in *D. radiodurans*

The *D. radiodurans* cells expressing DivIVA-RFP and FtsZ-GFP on plasmids were subjected to Z-stacking for 15-20 slices of 200 nm in each plane, and the localization of both the proteins with respect to each other was monitored. Several single cells were analyzed at different angles to understand the planes of both the proteins. Results shown here is a representative cell from the pool of identical cells (Fig 3). It showed that both the proteins get localized in different planes with respect to each other and the nucleiod. For instance, drDivIVA foci appeared in both conjunction and away from the nucleoid whereas FtsZ-foci appeared to be perpendicular to the nucleoid as shown by the x-y-z axis in the corresponding image (Fig 3A). The perpendicularity of Z-ring with respect to the genome is well known in rod-shaped bacteria. Thus, the drDivIVA protein localizes in two planes, (i) in the plane of FtsZ ring, and (ii) in the direction of genome segregation suggesting the possible role of this protein in both cell division and genome segregation by a mechanism unknown yet. The differential localization of both the proteins signifies that the FtsZ-ring form at the plane that is free from DivIVA thereby the plane of division might get determined in the coccus-shaped bacteria. In *B. subtilis*, DivIVA localizes at new division sites that would form new poles in the daughter cells (Eswaramoorthy et al., 2011). Interestingly, the T19A mutant appeared similar to that of wild type (Fig 3B). However, T19E mutant showed localization in a different pattern and the arrangement was not throughout the membrane rather at some predefined position and lost its dynamicity in the cells during growth (Fig 3C). These results suggested that RqkA phosphorylation of drDivIVA has affected the wild type function of this protein in both genome segregation and perpendicularity of cell division in *D. radiodurans*.

**Fig. 3:**
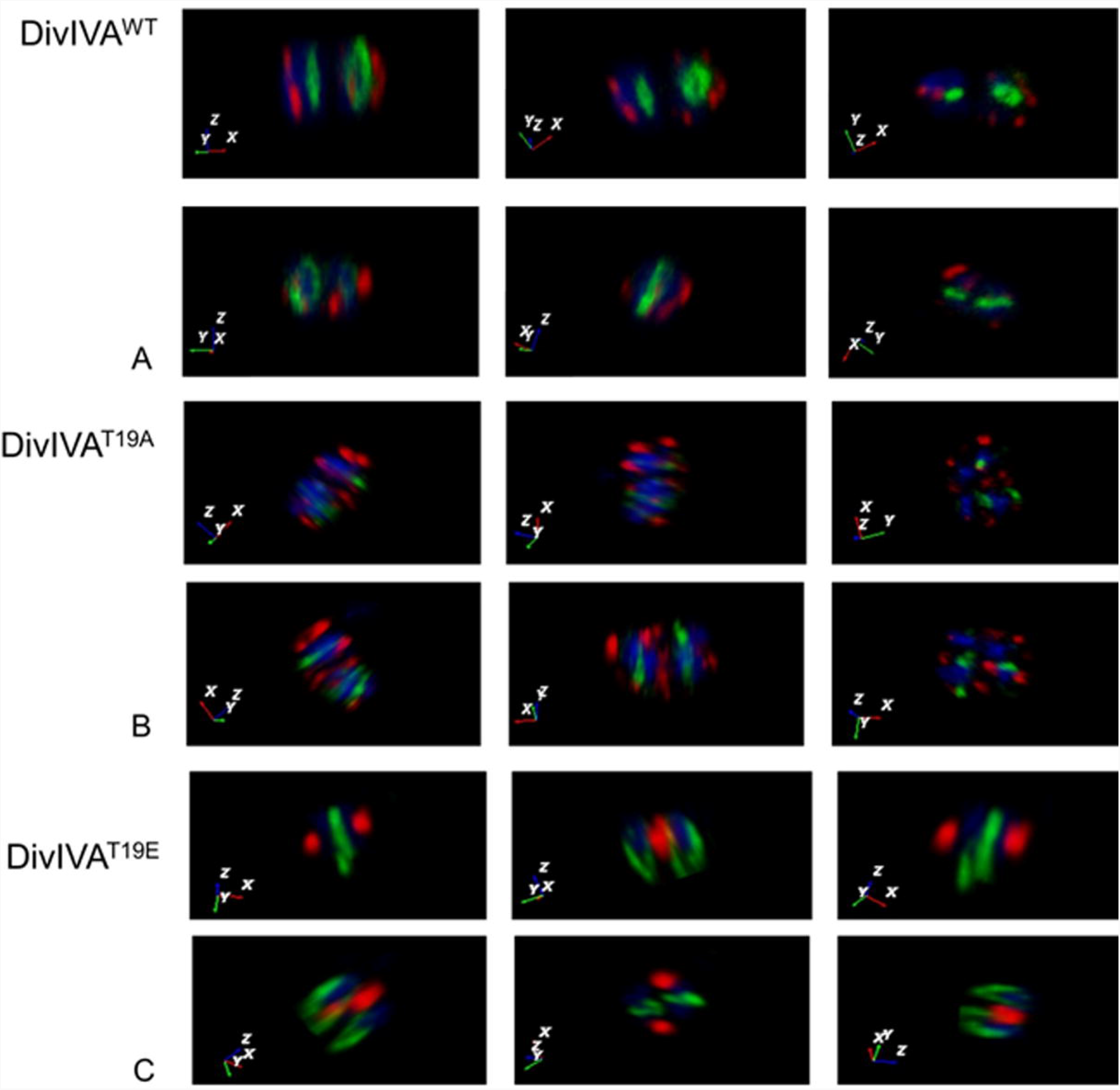
The localization of DivIVA/phospho-mutants and FtsZ in *D. radiodurans*. *D. radiodurans* expressing wild type DivIVA-RFP (DivIVA^WT^) and its phospho-mutants like DivIVA^T19A^ and DivIVA^T19E^ (red), FtsZ-GFP (green) and genome (blue) were analyzed for their planar position using confocal microscopy. The images were taken at several Z-planes and superimposed to give the overall 3-D picture and a single cell was analyzed by rotating at several angles and x-y-z axes are shown in the left bottom corner where they denote the position of DivIVA, FtsZ and genome respectively. The x-y-z axis is color coded according to the color of DivIVA (red), FtsZ (green) and nucleoid (blue). Data shown is a representative of a reproducible experiment repeated at least 3 times.

### Allelic replacement showed intolerance of phospho-mimetic allele under normal growth conditions

The *divIVA* is shown to be an essential gene for the normal growth of this bacterium, and its chromosomal copy could be replaced with selection marker only when drDivIVA was expressed episomally (Chaudhary et al., 2021). The effect of phosphorylation on drDivIVA functions was further evaluated by an allelic replacement of mutant copy with the wild type copy in the chromosome of this bacterium. The chromosomal copy of *divIVA* could be replaced with *divIVA-rfp* in *D. radiodurans* and the expression of DivIV-RFP under its native promoter was ascertained by fluorescence microscopy (Fig 4A). These cells showed normal growth and produced multiple DivIVA-RFP foci that were similar to the cells expressing DivIVA-RFP episomally *albeit* with a different intensity. For studying the cellular dynamics of T19A and T19E proteins when expressing under native promoter and their functional complementation, the replacement of wild type copy with *divIVA*^*T19A*^*-rfp* and *divIVA*^*T19E*^*-rfp* was attempted. The replacement of *divIVA* with *divIVA*^*T19A*^*-rfp* could be achieved and these cells showed normal growth and the wild type pattern of DivIVA^T19A^-RFP foci in the cells (Fig 4A). Notably, the homogeneous replacement of *divIVA* with *divIVA*^*T19E*^*-rfp* could not be achieved and the cells started debilitating upon approaching to homogeneity and finally ceased their growth under normal conditions. The colony-forming units (CFU) monitored during successive round of sub-culture showed a gradual decrease in prospective *divIVA*^*T19E*^*-rfp* cells (Fig 4B). However, this was achieved when drDivIVA was expressed *in trans* on plasmid. Interestingly, these cells grew slower in comparison with T19A and wild type controls (Fig 4C). The effect of DivIVA phosphorylation by StkP on the cell division has also been shown in *S. suis* (Ni et al., 2018). Surprisingly, the cells integrated with *divIVA*^*T19E*^*-rfp* allele in the presence of episomal drDivIVA also showed relatively less amount of DivIVA^T19E^-RFP protein, further indicating that phosphorylation seems to be deleterious for the stability of this protein in *D. radiodurans* (Fig 4D). The results presented here together suggested that phosphorylation of drDivIVA at T19 position seems making this protein non-functional/less stable and therefore, the *D. radiodurans* cells harbouring *divIVA*^*T19E*^*-rfp* in the place of *divIVA* could not be maintained under normal conditions.

**Fig. 4:**
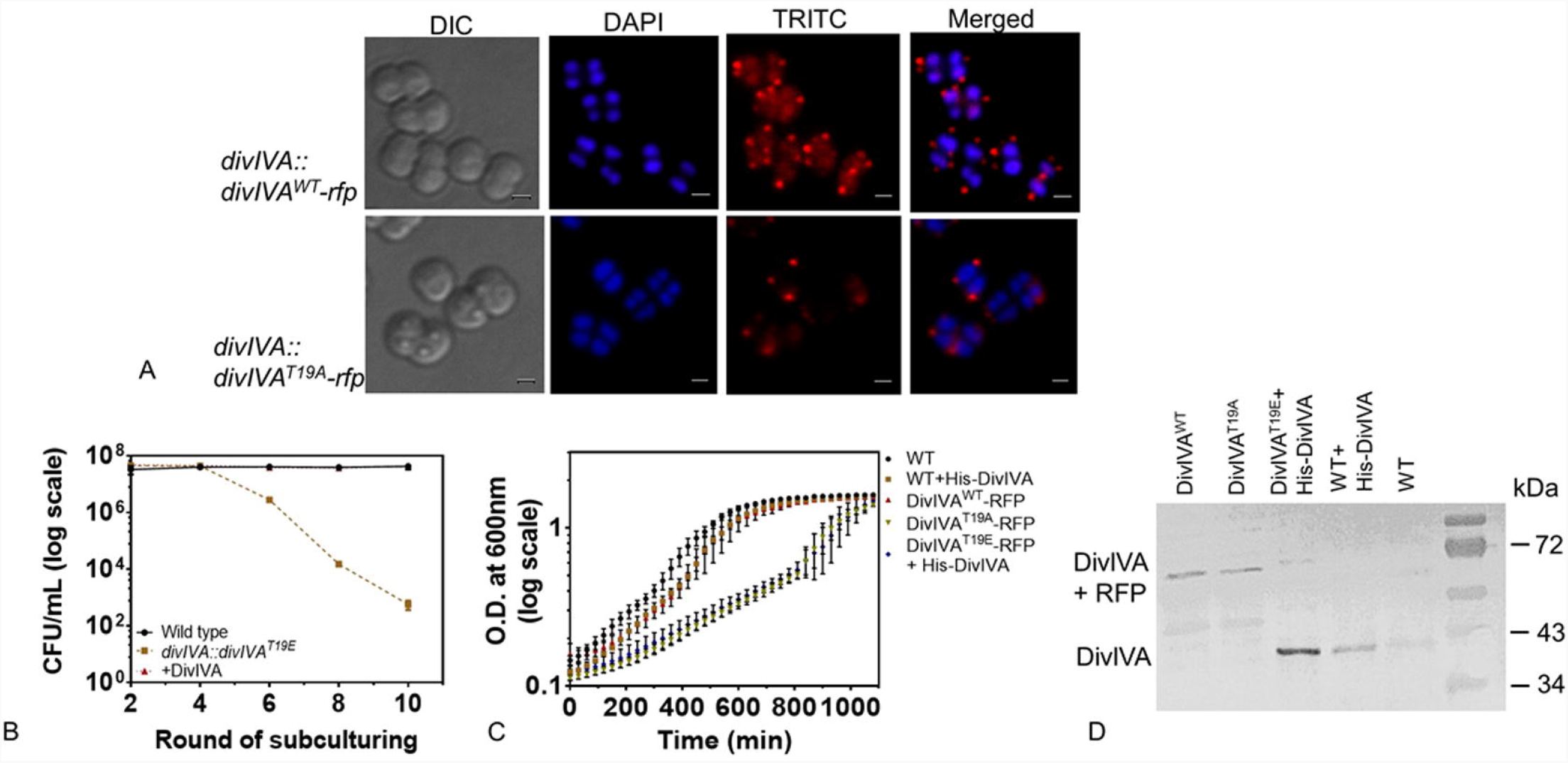
Chromosomal integration of *divIVA-rfp* variants under native promoter of *divIVA* in *D. radiodurans*. The divIVA coding sequences was replaced with the RFP fusion of wild type and phosphomutant of DivIVA in the chromosome of *D. radiodurans* as described in methods (Fig S2). The successful replacement of DivIVA-RFP and DivIVA^T19A^-RFP was confirmed by fluorescence microscopy **(A)**. The pNKD4A^T19E^ plasmid was linearized and transformed into *D. radiodurans* and transformants were scored in the presence of kanamycin. These cells were subcultured for several rounds in the presence of kanamycin. The possible replacement of *divIVA* with *divIVA*^*T19E*^*-rfp-nptII* was monitored on agar plate supplemented with antibiotic, after 1^st^, 3^rd^, 5^th^, 7^th^, 9^th^ and 11^th^ round of sub culture and compared with Δ*parA* mutant as the positive control. During these rounds of sub-culture, both wild type and prospective phospho-mimetic knock-in mutant of *divIVA* cells were platted on TGY and TGY + kanamycin (8µg/ml) agar plates, respectively. The colony-forming units (CFU) were obtained and plotted as a function of a round of sub-culture **(B)**. Growth-curve kinetics of wild type (WT), divIVA-rfp, (DivIVA^WT^-RFP) divIVAT19A-rfp (DivIVA^T19A^-RFP) and divIVAT19E expressing His-DivIVA episomally (DivIVA^T19E^-RFP + His-DivIVA) as well as wild type harbouring pRGhisD4A (WT + His-DivIVA) were monitored **(C)**. The total proteins of these knock-in derivatives of *D. radiodurans* were separated on SDS-PAGE and immunoblotted with anti-DivIVA antibodies **(D)**. Data shown without statistical attributes are the representatives of reproducible experiments repeated at least 3 times.

### drDivIVA marks the successive planes of cell division and regulates the position of FtsZ-ring formation

The cellular dynamics of drDivIVA (DivIVA-RFP) and FtsZ (FtsZ-GFP) was monitored in *D. radiodurans* growing under normal conditions. Time-lapse microscopic studies showed a cyclic change in the typical shapes of FtsZ-ring coinciding with different stages of cell cycle (Fig S2). For instance, the cells preparing for division have the closed form of Z-ring (Fig 5A time, t=0h), which changed to 3-shaped open form and finally returned to closed-form (t=3h) in cells producing two daughter cells (t=4h). Notably, the plane of opening of the FtsZ ring from close to 3-shaped was found to be perpendicular to the closed ring at t=0 h. To our surprise, DivIVA was localized in the mirror-image fashion in the two compartments of the tetrad. As the cell starts to divide, it would mark certain spatial territory and Z-ring formation would occur in the DivIVA free zone (t=1, 2h). When the cell division progressed towards the final stage, DivIVA spreads in its zone and the foci which were observed in the mother cell would have passed on daughter cells and may possibly serve as memory for the plane of previous division in the mother cells (black arrows). The daughter cells also possess DivIVA foci at the place where recent division has occurred (blue arrows; t=4h). These results showed that FtsZ ring is produced in DivIVA free space and the DivIVA moves from old division site to the new plane of cell division, and thus marks the site of FtsZ polymerization at juxtaposed position in the cells of this bacterium.

**Fig. 5:**
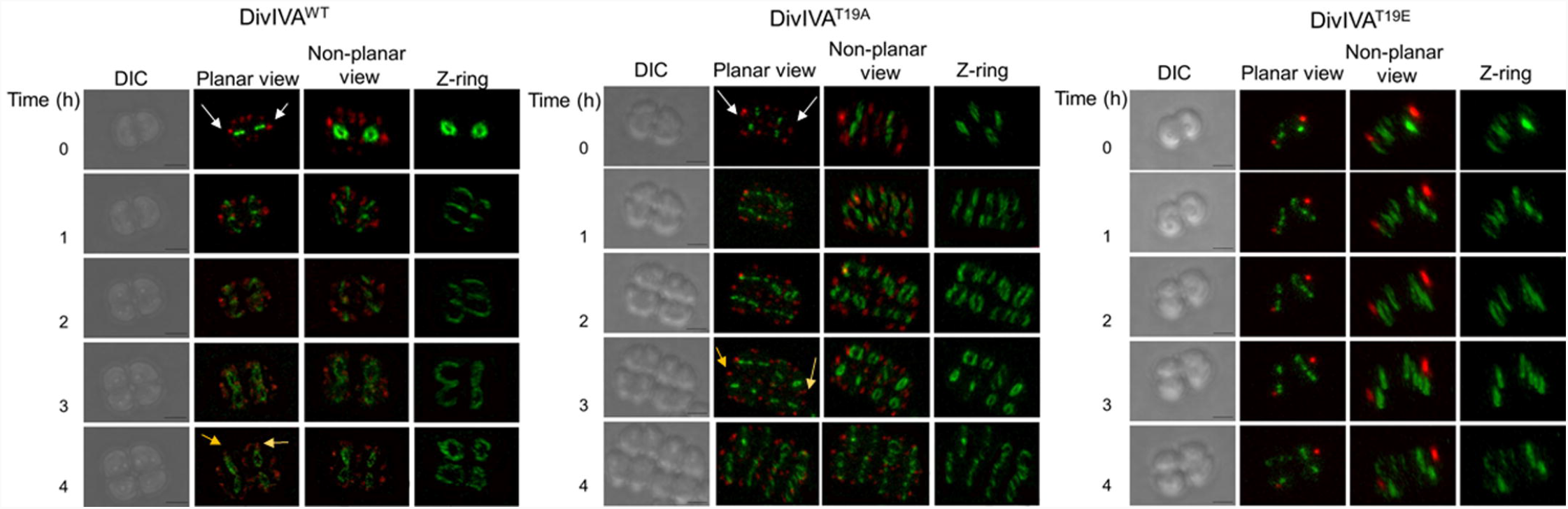
Time-lapse microscopy of the *D. radiodurans* cells expressing DivIVA-RFP derivatives and FtsZ-GFP. *D. radiodurans* cells expressing the RFP fusion of wild type DivIVA (A) DivIVA^T19A^ (B) and DivIVA^T19E^ (C) along with FtsZ-GFP were allowed to divide on microscopic slides and observed at different time points. Images were taken in DIC, GFP and RFP channels and presented as planner view as well as 3D views like non planner view and Z ring. The scale bars for all the imaging were kept constant to 500 nm. The cells proceeding towards the final stage of division and DivIVA observed in the mother cell have appeared in the daughter cells are marked with white arrow. A representative data of a reproducible experiment is shown.

### Unlike T19A and drDivIVA, the sub-cellular localization and dynamics of T19E were altered in *D. radiodurans*

Further the effect of drDivIVA phosphorylation on its localization and dynamics, FtsZ nucleation, and the dynamics of FtsZ ring polymorphism was also monitored in the cells expressing T19A and T19E mutants. The T19A mutant showed its movement to the new location with respect to the cell division plane and was found to be perpendicular to the plane of previous division which was very much similar with wild type protein (Fig 5, compare panels of DivIVA^WT^ and DivIVA^T19A^). However, the T19E mutants showed the contrasting phenotypes when compared with T19A and wild type DivIVA with respect to FtsZ-GFP conformation in *D. radiodurans* (Fig 5, DivIVA^T19E^). For instance, the opening of the Z-ring at the onset of division occurs in both the planes irrespective of the position of T19E and the cell division was completed but the foci of T19E protein did not move according to the cell division stage and only one fluorescent spot per cell of the dyad was observed as compared to multiple foci in case of DivIVA. These cells also showed cell cycle dependent FtsZ-ring polymorphisms as observed with wild type and T19A cells that could be attributed to the wild type background in respect of DivIVA. It may be noted that this study was carried out in wild type background because of technical inability (essentiality) to get chromosomal mutant of *divIVA*. The results obtained here clearly helped us to understand the effect of phosphorylation on the dynamics of DivIVA during normal growth of this bacterium. These results indicated that the T19E has failed to take part in cell division highlighting the importance of DivIVA phosphorylation in the regulation of cell division in this bacterium.

The significance of γ radiation inducible RqkA mediated phosphorylation on DivIVA dynamics was monitored in *rqkA* mutant exposed to γ radiation. The pattern of DivIVA localization has changed from multiple foci to less than four foci after 2 h of irradiation in wild type background whereas it remained unchanged in Δ*rqkA* background (Fig 6A). Both T19A and T19E mutants did not show any apparent change in their localization pattern after irradiation in both backgrounds. Interestingly, the localization pattern of DivIVA upon exposure with γ radiation had changed to nearly T19E as indicated by the lesser number of foci in comparison to multiple foci observed in case of T19A and DivIVA under normal conditions in the wild type background (Fig 6B). However, there was no change in the localization pattern of T19A and T19E proteins in both *rqkA* mutant and wild type backgrounds upon γ radiation exposure as well as DivIVA pattern in RqkA minus background. These results suggested that the γ radiation responsiveness of RqkA has affected DivIVA function in this bacterium.

**Fig. 6:**
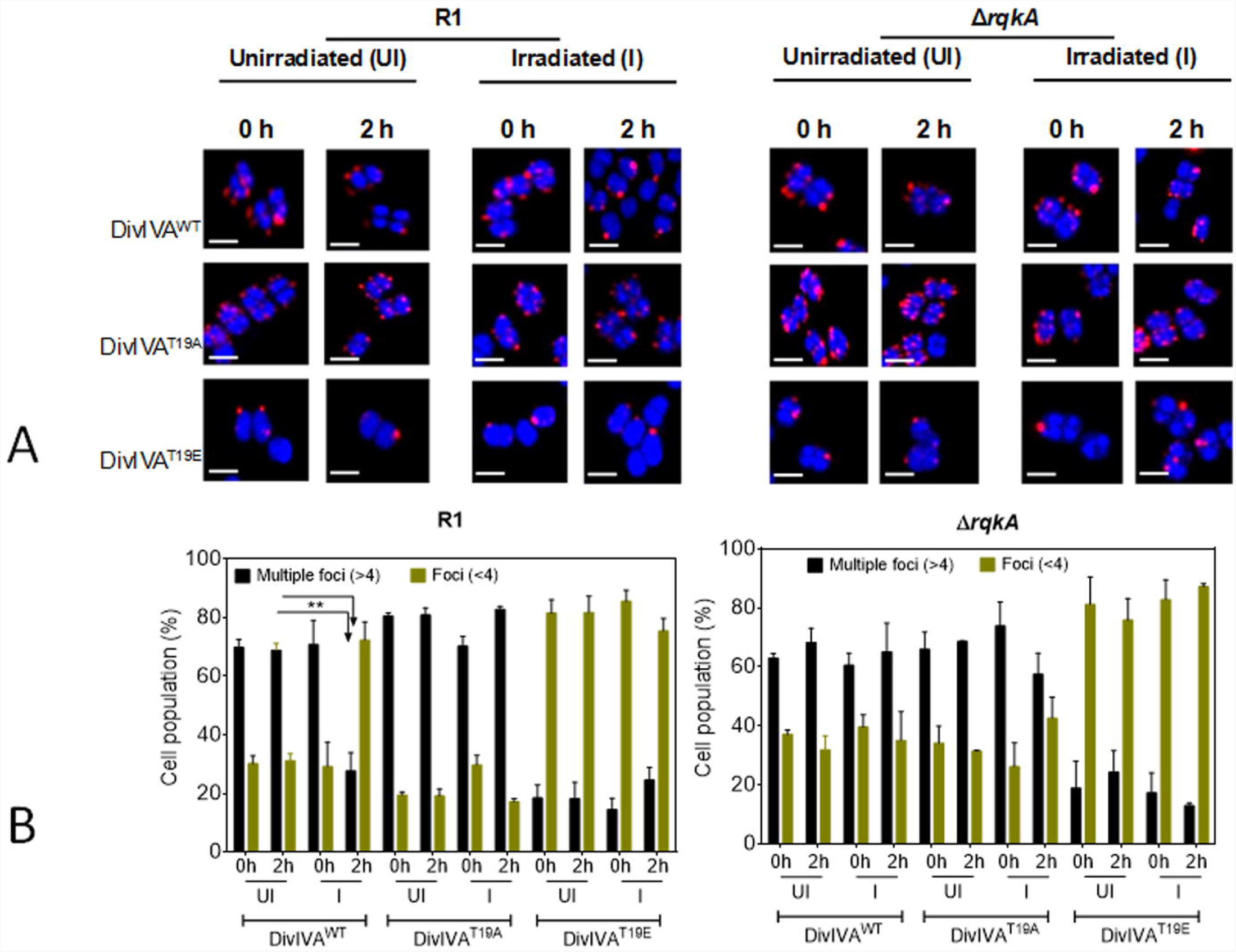
Localization pattern of DivIVA and its phosphomutants in the both wild type (R1) and Δ*rqkA* cells growth under unirradiated and irradiated conditions. The wild type (R1) and ΔrqkA cells expressing wild type (DivIVA^WT^), phosphoablative (DivIVA^T19A^) and phosphomimetic (DivIVA^T19E^) derivatives of DivIVA on plasmids pRGD4A, pRGD4A^T19A^ and pRGD4A^T19E^ respectively, were grown and exposed to 6 kGy γ-radiation (irradiated). A set of these cells were kept on ice as SHAM controls of irradiation (unirradiated). These cells were allowed for recover (PIR) as described in methods and cells at time 0PIR and 2hPIR were collected and processed for confocal microscopy at a scale bar of 1µm (A). Nearly 250 cells from each microscopic fields were taken and number of RFP foci per cell in these population were counted and a quantitative data is presented with statistical analysis (B).

### DivIVA seems to act as an anchor during chromosome segregation

The genome segregation is a pre-requisite for a true inheritance of genetic materials into daughter cells through binary fission in bacteria, and the interaction of DivIVA with its cognate genome segregation proteins have been reported (Chaudhary et al., 2019). Therefore, the genome segregation was monitored in cells expressing DivIVA variants by time-lapse microscopy. Earlier, five different morphologies; square, crescent, rod, branched and double rings, of nucleoid during genome segregation and their significance with different stages of cell growth have been suggested in *D. radiodurans* (Flóch et al., 2019). So, the mid-log phase cells expressing DivIVA-RFP was stained with Syto-green9 dye and the nucleoid morphology and movement was monitored. It was observed that these cells at t=0 have either square or early crescent shape nucleoid, which changes sequentially from crescent to rod to branched and finally double rings shaped during different stages of cell division and DivIVA showed dynamics along the movement of nucleoid (Fig 7). For instance, DivIVA is normally present at the cell periphery, which moves along the central septum in cells preparing for genome segregation (t=30). As the structure of nucleiod starts to change, DivIVA moves along the direction of genome movement, which might signify the DivIVA role in genome segregation. The DivIVA role in genome segregation has also been suggested in other bacteria (Hammond et al, 2019). Effect of T19 phosphorylation on genome segregation was less apparent possibly because of DivIVA heterogeneity by the expression of both wild type protein chromosomally and T19 mutants episomally. Although the pattern of T19E localization with respect to genome movement was same as wild type, a clear-cut arrest in its dynamicity was evidenced (Fig 7). The molecular basis of T19E effect(s) on genome segregation is not known yet. Earlier, drDivIVA interactions with cognate genome segregation proteins had been shown (Maurya et al 2018). So, the effect of phosphorylation’s mimic in the form of T19E on DivIVA interaction with ParA2 (ParA of chromosome II) protein was monitored *in vivo*. The drDivIVA and T19A proteins showed interaction with ParA2 while T19E protein did not interact with ParA2 *in vivo* (Fig 8) suggesting that phosphorylation of DivIVA at T19 position may have negatively affected its interaction with genome segregation proteins.

**Fig. 7:**
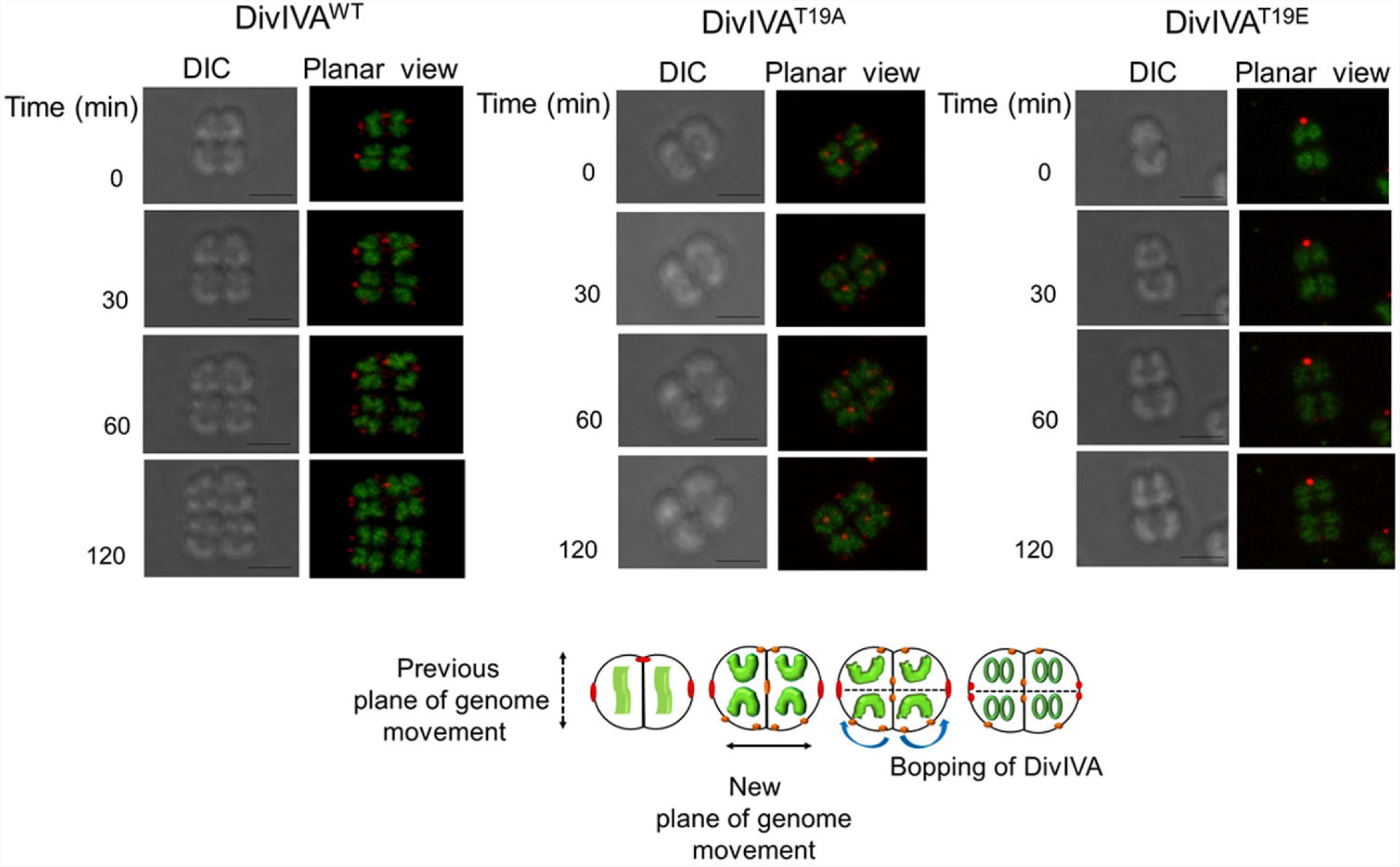
Time-lapse confocal microscopy of *D. radiodurans* expressing DivIVA-RFP with respect to genome movement during cell division. *D. radiodurans* cells expressing the RFP fusion of wild type DivIVA (A) DivIVA^T19A^ (B) and DivIVA^T19E^ (C) were stained for genome with Syto-green9 and allowed to divide cells on microscopic slides. These cells were imaged in DIC and RFP channels at different time points and images are presented in planner view. The scale bars for all the imaging were kept constant to 500 nm. A representative cell that exhibited duplicated copies of genome that changes during subsequent stages of cell division has been shown here for clarity (A). A general view of DivIVA and its mutant dynamics with respect to genome segregation as observed in majority of cell population is schematically represented for better clarity (B). Data shown is a representative of the reproducible experiments repeated 3 times.

**Fig. 8:**
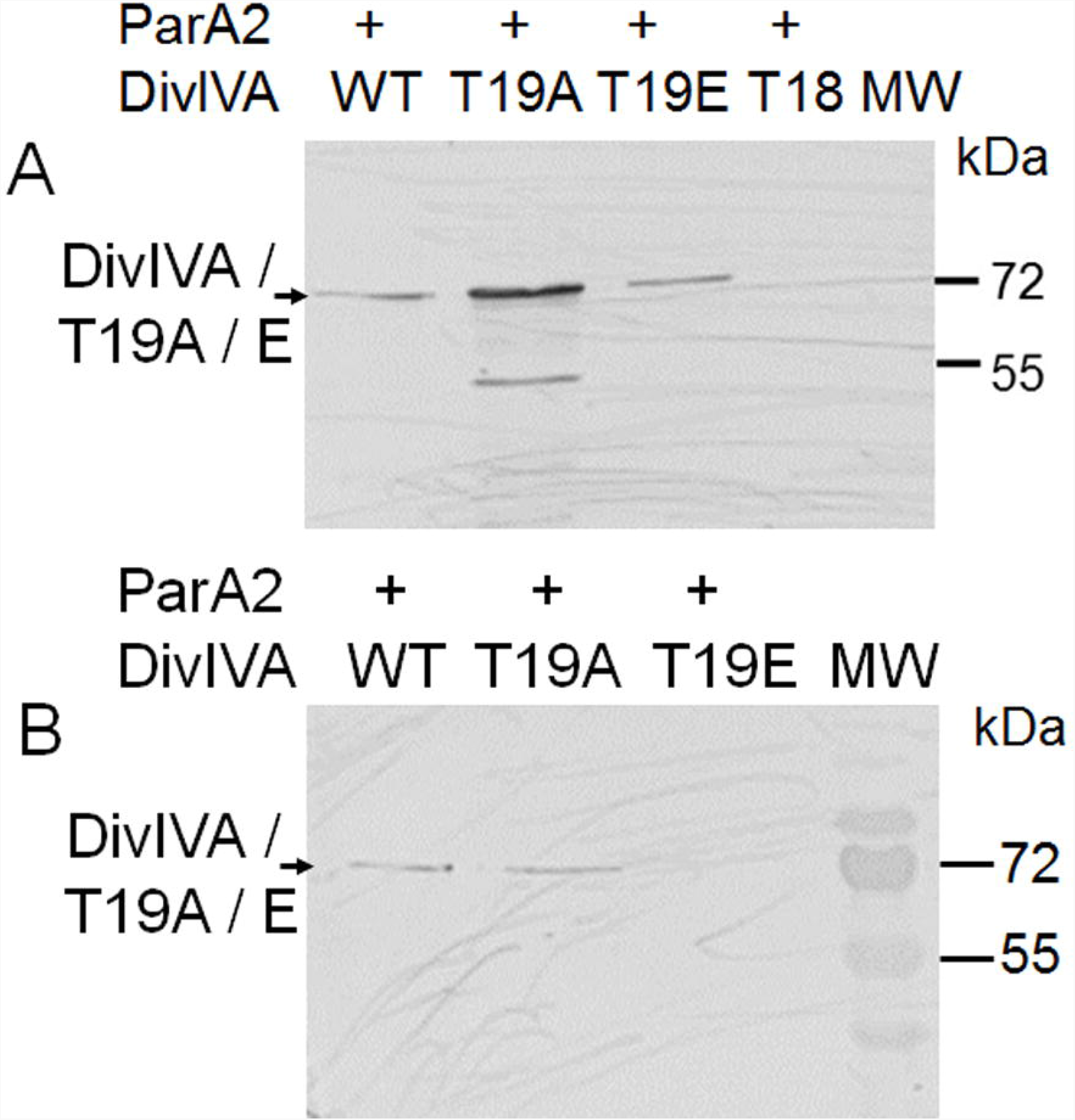
The effect of T19 replacement on the interaction with chromosome II ParA (ParA2) in vivo. Deinococcal cells co-harbouring pVA2T18 (expressing ParA2 with T18 tag) with pRGD4A (WT), pRGD4A^T19A^ (T19A) and pRGD4A^T19E^ (T19E) plasmid separately were checked for the expression of all three forms of DivIVA protein in cells. The cell lysates were run on 12% SDS-PAGE and immunoblotted with anti-DivIVA antibody (A). The cell free extracts from these cells were incubated with T18 antibody and immunoprecipitates were separated on SDS-PAGE and probed using DivIVA antibodies. Cells expressing T18 tag alone on pUT18 plasmid were used as negative control.

## Discussion

*Deinococcus radiodurans* a Gram-positive coccus bacterium, is characterized for its extraordinary resistance to DNA damage. The cytogenetic features of this bacterium include the presence of its majority population in tetrad form and a ploid multipartite genome system. Like other cocci, the plane of cell division is perpendicular to the previous plane of cell division. But unlike rod-shaped bacteria like *E. coli* and *B. subtilis*, there can be multiple virtual planes of cell division in cocci, and therefore the determination of mid-cell position for the initiation of cell division can become a deterministic factor of cell division in round-shaped bacteria. Further, *D. radiodurans* exposed to γ radiation shows growth-arrest till damaged genome gets repaired. This bacterium lacks LexA/RecA type canonical DNA damage response and cell cycle regulation (a paradigm in bacteria) and then, the molecular basis of growth arrest would be worth understanding. A DNA damage responsive Ser/Thr quinoprotein kinase (RqkA) has been characterized from this bacterium and its role in extreme phenotypes has been reported (Rajpurohit and Misra, 2010). RqkA phosphorylates both DNA repair and cell division proteins. Interestingly, the phosphorylation of DNA repair proteins enhances their activity while it negatively affects the functions of cell division protein FtsZ *in vitro* (Rajpurohit et al, 2022).

In *B. subtilis*, DivIVA localizes at both the poles and new division site (Edwards and Errington, 1997). DivIVAs have been found the substrates of eukaryotic like serine-threonine protein kinases (eSTKPs) in several bacteria including *S. pneumoniae, Mycobacterium spp*. The replacement of phosphorylation site of 201^st^ threonine with alanine in DivIVA has resulted into an elongated and bulged cell phenotype in *S. pneumoniae* (Fleurie et al., 2012). Further, the important roles of DivIVA in the determination of polarity and planes of cell division have been shown at least in cocci (Chaudhary et al., 2021). DivIVA in general localises at the negative curvature, which is also marked by cardiolipin microdomains at least in *B. subtilis* (Renner and Weibel, 2011). The presence of DivIVA at new division sites has also been shown in *S. aureus* and *S. pneumoniae* (Pinho and Errington, 2004) that also follows the alternate plane of septal growth during the cell division. It has been shown that MapZ protein that determines the FtsZ positioning in the membrane of *S. pneumonia*e is conserved in *streptococci, lactococci* and most *enterococci* (Fleurie et al., 2014). Further, the interaction of its both N-terminal and C-terminal sub domains is mandatory for its function of placing cognate FtsZ in mid-cell position for septal growth (Manuse et al., 2016). The site-directed mutagenesis in Wag31 revealed that phosphorylation regulates the activity of this protein in mycobacteria (Kang et al., 2008).

*D. radiodurans* lacks cardiolipin microdomains but its DivIVA (i) undergoes phosphorylation by RqkA at T19 site that maps in proximity with the polar determining motif, (ii) contains an extended CTD when compared with DivIVA of other bacteria, (iii) is an essential protein and its CTD domain contributes to the positioning of septum during normal growth in this bacterium (Chaudhary et al, 2021), and (iv) interacts with cognate cell division and genome segregation proteins (Maurya et al, 2018). Therefore, the localization of DivIVA and its phosphomutants, and their dynamics with respect to FtsZ ring positioning and genome segregation was monitored. It was observed that at the initiation of cell division, the wild type DivIVA (mostly dephosphorylated form) localizes at cell membrane, marks spatial territories for the Z-ring formation to take place in the zone that is free from DivIVA (Fig 9). During next cycle of cell division, DivIVA spreads in its zone and the foci, which were observed in the mother cells have now passed on to daughter cell, possibly acting as the memory foci for cell to determine the plane of next cell division that would be perpendicular to previous plane of division. The daughter cells also possess foci at the place where new cell division would occur. Thus, it is observed that DivIVA marks both old and new planes of cell division in this coccus bacterium. When the effect of T19 phosphorylation on this characteristic of DivIVA was investigated, the T19A behaved similar to wild type. However, both localization and dynamics of T19E mutant during cell division were found to be different from wild type and T19A mutant.

**Fig. 9:**
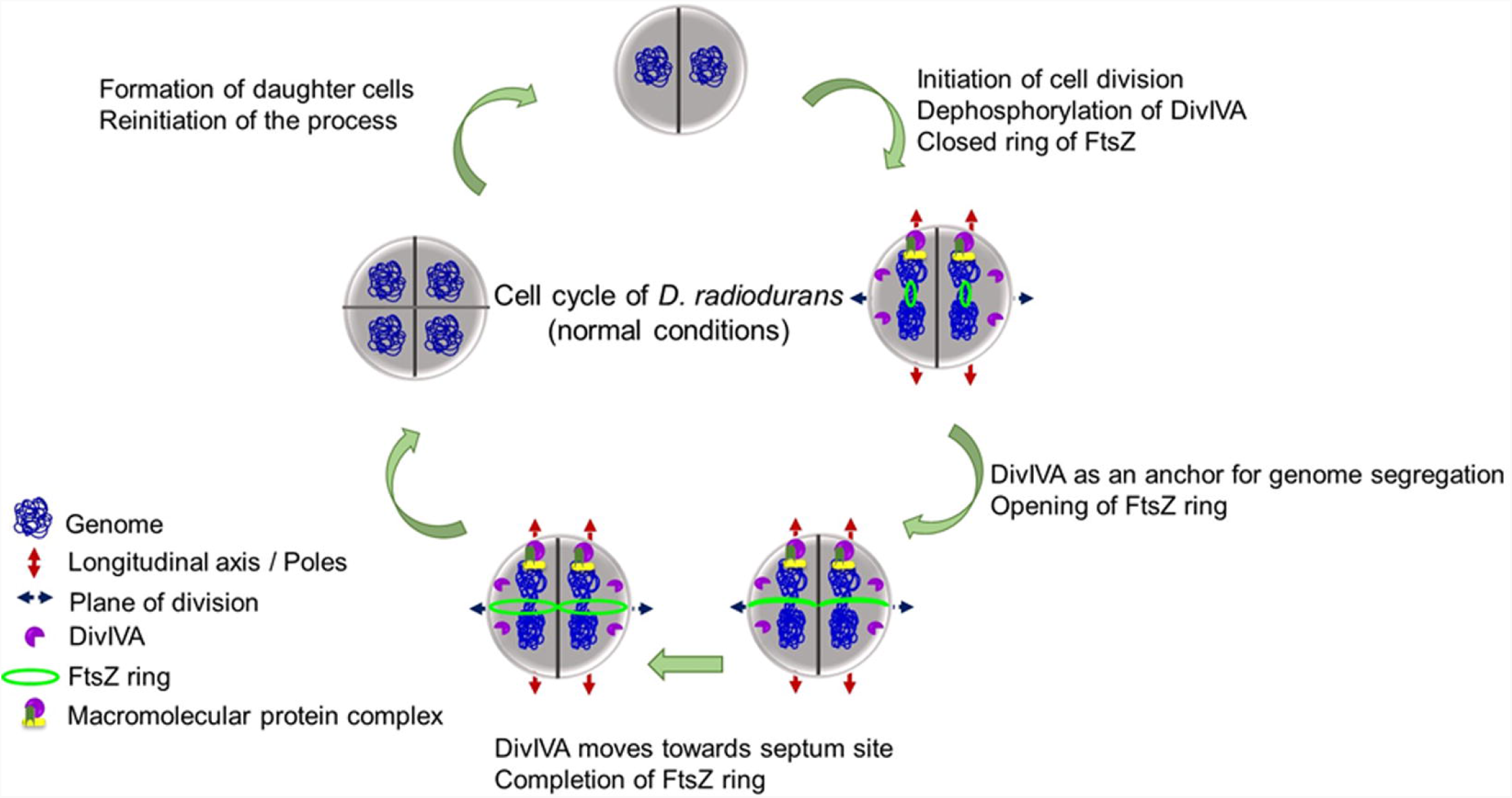
A model depicting steps associated with cell cycle of *D. radiodurans* under normal conditions. Schematic representation of the findings from this study that mostly fit into a model of cell division and genome segregation during normal growth of *D. radiodurans*.

Time-lapse microscopic studies clearly suggested that T19E does not show dynamics during the cell division suggesting a strong possibility of phosphorylation mediated arrest on DivIVA dynamics in this bacterium. A distinct effect of γ radiation on the dynamics of wild type DivIVA in the presence and absence of RqkA (wild type and *rqkA* mutant, respectively) while nearly no change in the localization pattern and dynamics of phospho-mutants in these genetic backgrounds strongly supported the effect of T19 phosphorylation by RqkA on DivIVA functions in *D. radiodurans*. Although, the molecular mechanisms explaining the loss of DivIVA dynamics upon T19 phosphorylation would be worth studying independently, the available results presented here suggest the DivIVA phosphorylation at T19E site by RqkA and this phosphorylation of DivIVA helps the bacterium to arrest its growth in response to γ radiation damage as the mechanism of cell division regulation in *D. radiodurans*. Furthermore, DivIVA roles in the determination of both cell polarity and alternate planes of cell division as well as if it contributes to cell-cycle regulation in response to γ radiation could have not been shown in any other bacteria but in *D. radiodurans* that shows resistant to DNA damage. Therefore, this study signifies not only with a demonstration of DivIVA phosphorylation by any STPK but by a radiation responsive STPK (RqkA) and this event could lead to cell cycle arrest in this bacterium.

## Materials and Methods

### Bacterial strains, plasmids and bacterial growth measurement

*D. radiodurans* R1 (ATCC13939) was a kind gift from Prof. J Ortner, Germany (Schäfer et al., 2000). Shuttle plasmids pVHS559 and pRADgro were suitably modified in our laboratory from the parental backbone of p11559 (Lecointe et al., 2004) and pRAD1 (Meima and Lidstrom, 1996), respectively. The *E. coli* cells containing pVHS559 and its derivatives were maintained in the presence of spectinomycin (40 µg/ml) while *D. radiodurans* was maintained in the presence of spectinomycin (75 µg/ml) as described (Charaka et al., 2012). Similarly, pRADgro plasmid was maintained in *E. coli* with ampicillin (100 µg / ml) and in *D. radiodurans* with chloramphenicol (8 µg / ml) (Misra et al., 2006). Details of the plasmids and strains used in this study are listed in Table 1. The derived strains were grown in the presence of respective antibiotics and their growth at OD_600_ was monitored in sterile 24-well microtitre plate using Synergy H1 Hybrid multi-mode microplate reader, Biotek for overnight at 32°C. The data was processed, analyzed for statistical significance and plotted using GraphPad Prism software.

**Table 1:**
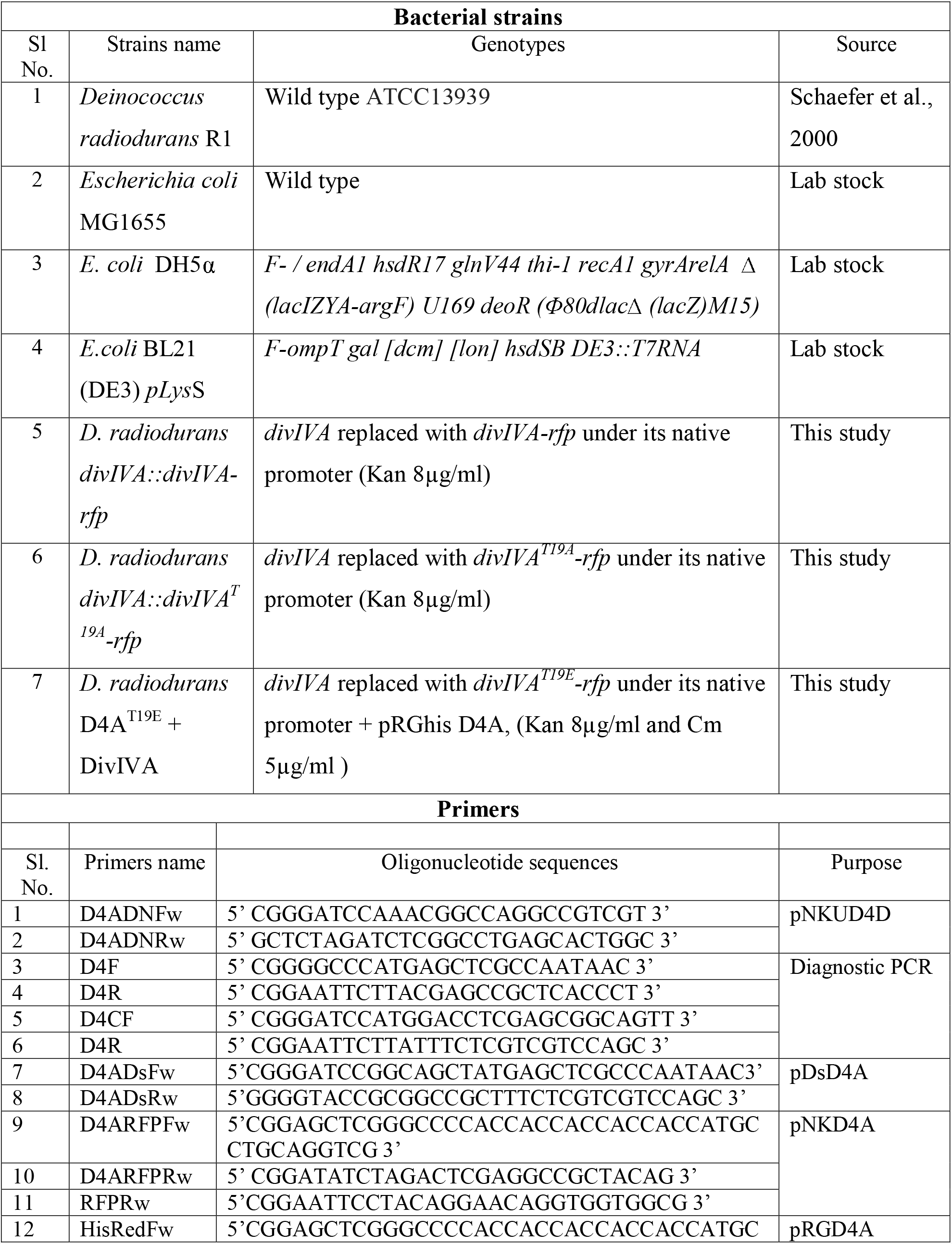
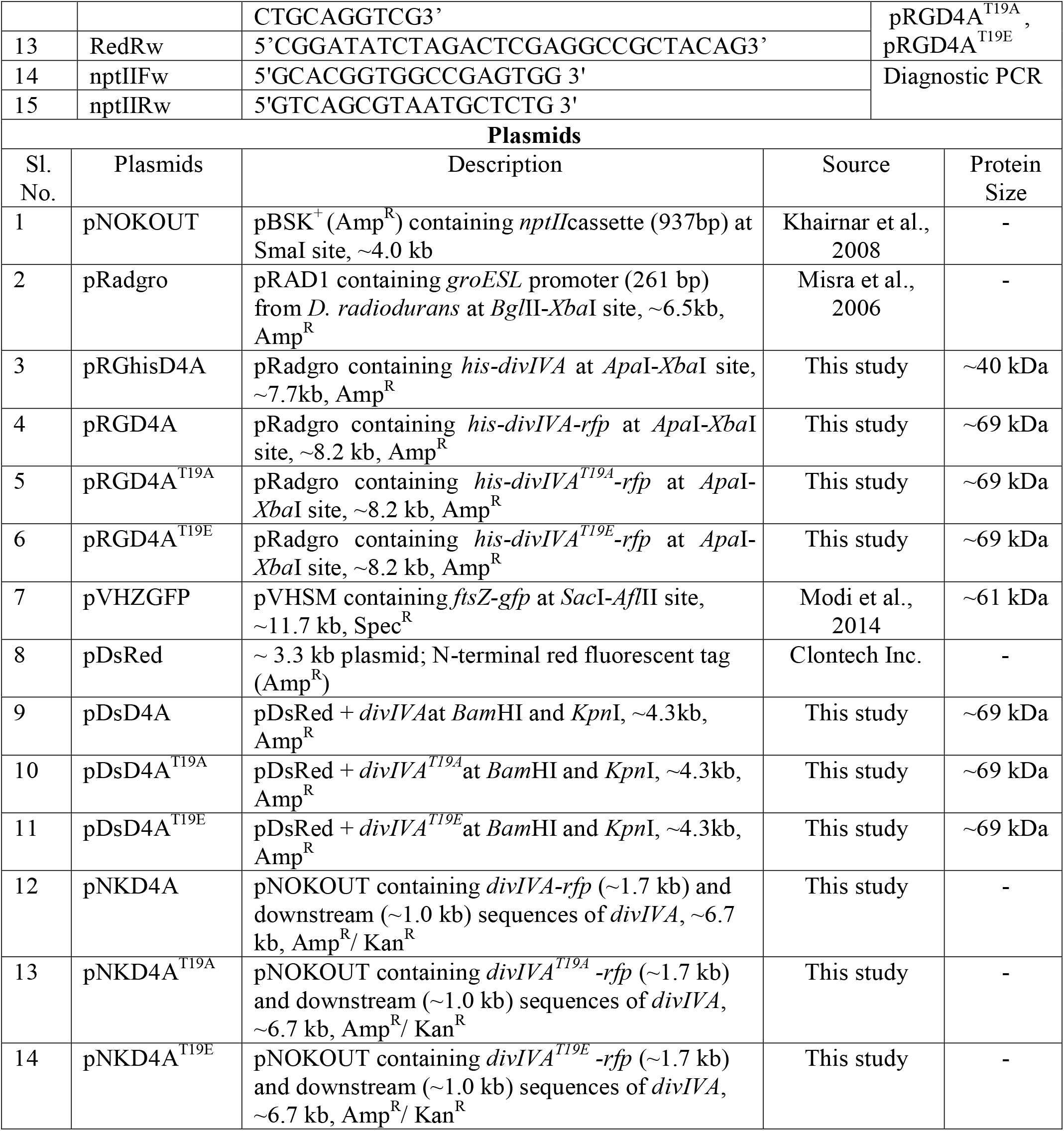
List of strains, primers and plasmids used in the study.

### Protein phosphorylation and immunoblotting

The recombinant drDivIVA and RqkA were expressed on pETD4A (Chaudhary et al, 2019) and pET2518 (Rajpurohit and Misra, 2010) plasmids, respectively in transgenic *E. coli* BL21 and purified using protocols as described earlier. The recombinant DivIVA was incubated in different molar ratios (X/100, X/25, X/10, where X is DivIVA) of recombinant RqkA in the kinase reaction mixture containing 10 mM Tris-HCl (pH 7.6), 20 mM KCl, 1.5 mM MgCl_2_, 0.5 mM dithiothreitol, 2% glycerol, 0.1 mM EDTA, and 5 mM cold-ATP and [γ-^32^P] ATP, for 30 min at 37°C. The reaction was stopped by adding 2× SDS sample buffer and the phosphorylation was checked on 12% SDS-PAGE as described earlier (Maurya et al., 2018). The gel was dried and exposed to X-ray film and autoradiograms were developed. The data was further analyzed and plotted using Prizm software.

The phosphorylation of DivIVA by RqkA was further checked *ex-vivo* in *E. coli* BL21 cells co-expressing drDivIVA and RqkA episomally. The co-expression of both the proteins was ascertained independently using specific antibodies. The recombinant drDivIVA protein was purified from the *E. coli* cells co-expressing RqkA on plasmid and checked for phosphorylation using phospho-Ser/Thr epitopes antibodies as described earlier (Rajpurohit and Misra, 2010). In brief, purified drDivIVA from *E. coli* expressing RqkA (P-DivIVA) and control (DivIVA) were loaded on 12% SDS-PAGE and transferred to the PVDF membrane. Blot was hybridized with polyclonal phospho-Ser/Thr epitopes antibodies and signals were detected using AP conjugated anti-rabbit secondary antibody. Similarly, the phosphorylation status of DivIVA from *D. radiodurans* (both unirradiated (UI) and irradiated (I)) was checked by immunoblotting. The cell lysates of *D. radiodurans* expressing pRGhisD4A were loaded on 12% SDS-PAGE and transferred to the PVDF membrane and blot was hybridized with both anti-DivIVA and polyclonal phospho-Ser/Thr epitopes antibodies.

### Mapping of phosphorylation sites in drDivIVA

drDivIVA co-expressing with recombinant RqkA in *E. coli* and control was purified to homogeneity. Proteins were separated on SDS-PAGE and stained with Coomassie brilliant blue. The DivIVA protein band was incised from the gel and reduced with 5 mM TCEP followed by alkylation with 50 mM iodoacetamide before it was digested with 1µg trypsin at 37°C for 16 h. Peptide mixture was purified using C18 column (The Nest Group, Southborough, MA) according to the manufacturer’s protocol and dried under vacuum. The peptide pellet was dissolved in buffer A (5% acetonitrile / 0.1% formic acid) and mixture was resolved using 15cm PicoFrit column (360 um OD, 75 um ID, 10 um tip) filled with 1.8 μm C18-resin (Dr. Maeisch) in EASY-nLC 1000 system (Thermo Fisher Scientific) coupled to QExactive mass spectrometer (Thermo Fisher Scientific), by outsourcing. In brief, the peptides were loaded with buffer A and eluted with a 0-40% gradient of Buffer B (95% acetonitrile/0.1% formic acid) at a flow rate of 300 nl/min for 40 min. The QExactive was operated using the Top10 HCD data-dependent acquisition mode with a full scan resolution of 70,000 at m/z 400. MS/MS scans were acquired at a resolution of 17500 at m/z 400. Lock mass option was enabled for polydimethylcyclosiloxane (PCM) ions (m/z = 445.120025) for internal recalibration during the run. The data was processed using proteome discoverer software against Uniprot *D. radiodurans* database with the phosphorylation as dynamic modification criteria.

### Site-directed mutagenesis

Site directed mutagenesis was carried out using Q5® Site-Directed Mutagenesis Kit using manufacturer’s protocol (source of kit, NEB or else) using primers carrying the altered codon were designed using the online tool OligoCalc. In brief, the primers containing desired mutation in threonine codon so as to change into either alanine or aspartate codon was used for amplification of the plasmid carrying *divIVA* (pUTD4A) using standard procedure. PCR products were incubated with enzyme mix containing a kinase, a ligase and *Dpn*I at 25°C for 30 min. Together, these enzymes allow for the rapid circularization of the PCR product and removal of the non-mutated DNA template by *Dpn*I. The circularized plasmid was transformed into *E. coli* NovaBlue cells and mutations were ascertained by sequencing. The mutated alleles expressing T19A and T19E mutant version of DivIVA were cloned separately in different expression plasmids.

### Construction of recombinant plasmids

The recombinant plasmids pDsD4A and pRGD4A were constructed for the monitoring of the localisation of DivIVA (DR_1369) as DivIVA-RFP. Similarly, pDsD4A^T19A^ and pRGD4A^T19A^, and pDsD4A^T19E^ and pRGD4A^T19E^ plasmids were constructed for expression of T19A-RFP and T19E-RFP, respectively. In brief, the coding sequences of DivIVA and its T19A and T19E mutant alleles were PCR amplified using sequence-specific primers (Table 1) and cloned at *Bam*HI-*Kpn*I sites in pDsRed plasmid (Clontech Inc.) yielding pDsD4A and pDsD4A^T19A^, pDsD4A^T19E^, respectively. Further, the translation fusion of *divIVA-rfp, divIVA*^*T19A*^*-rfp* and *divIVA*^T19E^-*rfp* was amplified from pDsD4A, pDsD4A^T19A^and pDsD4A^T19E^, and respectively using RedFw and RedRw primers and cloned in pRadgro to yield pRGD4A, pRGD4A^T19A^ and pRGD4A^T19E^, respectively. The recombinant plasmids pNKD4A, pNKD4A^T19A^ and pNKD4A^T19E^ were constructed to replace the wild type *divIVA* allele with *divIVA-RFP, divIVA*^*T19A*^*-rfp* and *divIVA*^T19E^-*rfp* alleles in the chromosome for the expression of DivIVA-RFP, T19A-RFP and T19E-RFP, respectively under the native promoter in *D. radiodurans*. For that, *divIVA-rfp* alleles and downstream region of the coding sequence of DivIVA were separately PCR amplified using sequence-specific primers (Table 1). The *divIVA-rfp* and its mutant alleles were cloned at *Apa*I-*Eco*RV while the downstream sequences of DivIVA at *Bam*HI-*Xba*I sites in pNOKOUT (Kan^R^) to give pNKD4A, pNKD4A^T19A^ and pNKD4A^T19E^, respectively.

### Generation of chromosomal insertions in *divIVA*

For generating chromosomal insertions, the *D. radiodurans* cells were transformed with *XmnI* linearized pNKD4A, pNKD4A^T19A^ and pNKD4A^T19E^ plasmids using protocols as described in (Khairnar et al., 2008). The transformants were grown for several rounds in the presence of kanamycin (8μg/ml). The cycles of alternate streaking on TYG agar plate and sub-culturing in TYG broth were repeated for several rounds till homogenous replacement of target with insertion sequences has been achieved (Chaudhary. et al., 2021). Genomic DNA was prepared and homogenous replacement of *divIVA* with the expressing cassette of *divIVA-rfp-nptII* through homologous recombination was ascertained by diagnostic PCR (Fig S3) with gene specific primers as given in Table 1. The PCR products were analysed in 1 % agarose gel. The replacement of wild type allele with T19E allele could not be obtained as the cells did not survive. For creating *divIVA*^T19E^-*rfp* insertion in place of *divIVA* allele was achieved when DivIVA was expressed episomally on pRGhisD4A plasmid. The confirmation of insertions of *rfp-nptII* downstream to *divIVA* in the correct reading frame was monitored as the expression of DivIVA–RFP and its variants under native promoter, by fluorescence microscopy.

### Fluorescence microscopy and image analysis

General procedures of fluorescence microscopy using confocal microscope (Model Olympus Cell IX 83 Fluoview 3000) were as described earlier (Chaudhary et al, 2021). In brief, the bacterial cultures were grown in TYG broth, fixed with 4% paraformaldehyde for 10 min on ice and washed two times with phosphate-buffered saline (pH 7.4). These cells were stained with DAPI (0.5 µg/µl) for 10 min on ice and then washed three times with PBS. The cells were resuspended in PBS, mounted on 1% agarose bed on glass-slides and samples were observed under microscope. For FtsZ localization, the cells expressing FtsZ-GFP on pVHZGFP plasmid (Modi et al., 2014) were grown overnight and cells equivalent to 0.05-0.1 OD_600_ were diluted with a fresh medium containing required antibiotic(s) and induced with 10 mM IPTG overnight. For time-lapse imaging, the exponentially growing cells were suspended in PBS and mounted on an agarose pad made in 2XTGY and designed with air holes to oxygenate the cells. Confocal microscopy was performed with the laser beams focused to the back focal plane of a 100 × 1.40 NA oil-immersion apochromatic objective lens (Olympus). The laser parameters for illumination at the sample were tuned using the installed FLUOVIEW software. For imaging at required time points, the series of Z-planes were acquired every 400-nm using the motorized platform. Fluorescence emission was collected through a DM-405/488/561 dichroic mirror and the corresponding single-band emission filters. For constructing 3D images, the cells expressing FtsZ-GFP and DivIVA-RFP or its phospho-mutants were sliced in Z planes at every 400 nm at 45-60 min interval for a period of 4-5 h using very low powers of lasers at 561 nm and 488 nm. For monitoring the movement of the genome, the cells expressing DivIVA-RFP and its phospho-mutants were stained with 150nM of Syto-green9 dye and images were acquired for a period of 2-3 h after every 30-45 min using very low powers of lasers at 561 nm and 488 nm. For image analysis, cells were taken from at least two separate microscopic fields of the images captured in two independent experiments and analyzed for required attributes. The image analysis and other cell parameters were measured using automated Cell Sens software. Data obtained were subjected to Student’s t-test analysis using statistical programs of Graphpad Prism 5.0.

For imaging of DivIVA and its phopho-mutant forms in the both wild type and Δ*rqkA* mutant background, the cells in exponential phase were exposed to 6 kGy dose of γ radiation whereas control cells were kept on ice. After γ-irradiation, both un-irradiated (UI) and irradiated (I) cells expressing DivIVA^WT^, DivIVA^T19A^ and DivIVA^T19E^ were collected at 0h and 2h of post-irradiation recovery. The cells were processed for confocal microscopy as described above. Images were processed using Cells Sens software and Adobe Photoshop 7.0.

### Co-immunoprecipitation (Co-IP) of drDivIVA and its phospho mutant forms with ParA2

*D. radiodurans* R1 cells expressing DivIVA-RFP (wild type and phospho-mutant forms) were transformed with ParA2-T18 (pVT18A2; Maurya et al., 2019) and colonies were scored on spectinomycin (70 µg/mL). These colonies were further grown and induced with 20 mM IPTG overnight. The cell-free extracts of *D. radiodurans* expressing these constructs along with vector control were prepared. In brief, deinococcal cells expressing the desired proteins were pelleted and washed with 70% ethanol followed by a wash of phosphate buffer saline. Pellets were suspended in 500 µl of lysis buffer A (50 mM Tris, pH 7.5, 100 mM NaCl, 1 mM PMSF, 5 mM MgCl_2_, 1 mM dithiothreitol [DTT], 0.5 % Triton X-100) with 0.5 mg/ml Lysozyme and 50 μg protease inhibitor cocktail tablet (Roche Biochemicals) followed by sonication on ice. Cell debris was removed by centrifugation at 2,000 × g for 10 min at 4°C. The clear cell-free extracts were immunoprecipitated using monoclonal antibodies against T18 tag following Protein G Immunoprecipitation Kit (Merck Inc) protocol. Immunoprecipitates were separated on 10% SDS-PAGE and blotted onto PVDF membrane and hybridized with anti-DivIVA antibody. Hybridization signals were detected using anti-mouse secondary antibodies conjugated with alkaline phosphatase using BCIP/NBT substrates (Merck Inc.).

## Data Availability

This study includes no data deposited in external repositories

## Acknowledgements

Authors would like to thank Dr. Sebastian Raja for his initial demonstration of the confocal microscope and Dr. Ganesh K Maurya for his critical reading of the manuscript and comments. Reema Chaudhary is grateful to Department of Atomic Energy, India for her fellowship.

## Author contributions

**RC**- Experiments, data analysis, discussion and manuscript writing

**SK**- Experiments, data analysis, discussion and manuscript writing

**HSM**- PI, conceptualization of idea, data analysis, discussion, manuscript writing and communication for publication.

## Declaration

This work has been a part of doctoral thesis of Dr. Reema Chaudhary, submitted to Homi Bhabha National Institute (Department of Atomic Energy-Deemed to be University), Mumbai.

## Supplementary figures legend

**Fig. S1: Phospho-site mapping of deinococcal DivIVA.** The phosphorylated DivIVA protein was subjected to mass-spectrometry for identification of the phospho-site. A chromatogram of the protein was generated and ion-series of the phospho-peptide fragment were also analyzed with respect to non-phospho form of the protein **(A-B)**. Pair-wise alignment of the first few amino acids of DivIVA of *B. subtilis* and *D. radiodurans* was highlighted to show the proximity of phosphorylation and polar motifs **(C)**. Phospho-peptide was found to be DI**\*T\***HQSFDGR where T is mutated to alanine (A; phospho-ablative) and glutamate (E; phospho-mimetic) by site-directed mutagenesis **(D)**. The mutant forms were also checked for phosphorylation *ex-vivo. E. coli* cells harbouring pRadRqkA and any one of the following plasmids, pRGD4A (DivIVA^WT^), pRGD4A^T19A^ (DivIVA^T19A^) and pRGD4A^T19E^ (DivIVA^T19E^) were grown. The samples were run on 12% SDS-PAGE and immunoblotted with anti-phospho Ser/Thr antibody **(E)**. *E. coli* harbouring pRadRqkA vector was used as a control.

**Fig. S2: Time-lapse confocal microscopy of *D. radiodurans* expressing DivIVA-and DivIVA^T19A^-RFP from its native promoter and FtsZ-GFP.** The *D. radiodurans* cells co-expressing DivIVA-RFP (A) or DivIVA^T19A^-RFP (B) and FtsZ-GFP was processed for time-lapse microscopy and cells were imaged for 3-4 h for every 1 h with the scale bar of 500 nm. The signal of DivIVA-RFP has bleached after 1 h so different cells were taken from the field at 0 and 1 h of time point to represent the dynamics of native DivIVA-RFP and stages like closed, 3-shaped and intermediate stage of FtsZ-GFP ring during the cell division (C).

**Fig. S3: A diagrammatic representation of the strategy employed for replacement of divIVA coding sequences with RFP fusions of wild type and mutant alleles in the chromosome of *D. radioduarns*.** The *divIVA-rfp* and downstream sequences of *divIVA* were cloned into pNOKOUT vector yielding pNKD4A **(A)**. The same strategy was followed for creating other mutant constructs where *divIVA*^*T19A*^*-rfp* and *divIVA*^*T19E*^*-rfp* were cloned at upstream of *nptII* in pNOKOUT yielding pNKD4A^T19A^ and pNKD4A^T19E,^ respectively (A). The plasmids was linearized and transformed into *D. radiodurans* cells and transformants were scored in the presence of kanamycin. The integration of *rfp* downstream to *divIVA* followed by antibiotic resistance marker gene (nptII) was confirmed by diagnostic PCR using primers as described in Table 1 **(B)**. The PCR products in the gel are labelled for their respective identity as following, *nptII* (+) for antibiotic resistance gene from pNOKOUT; *nptII* from the corresponding knock-in strain; *divIVA* (from R1), *divIVA*^*WT*^ (from *divIVA::divIVA-rfp), divIVA*^*T19A*^ (from *divIVA::divIVA*^*T19A*^*-rfp)* and *divIVA*^*T19E*^ (from D4A^T19E^ + DivIVA); *divIVA*^*DN*^ for downstream sequence of *divIVA; divIVA*^*WT*^ –*rfp, divIVA*^*T19A*^-*rfp* and *divIVA*^*T19E*^-*rfp* for *divIVA* along with *<u>rfp</u>* in all three types of strains; I indicates the product of *divIVA-rfp-nptII-divIVA*^*DN*^ from the corresponding strains.

