## Supplementary figures and images for "DivIVA phosphorylation at threonine affects its dynamics and cell cycle in *Deinococcus radiodurans*"

### Fig S1

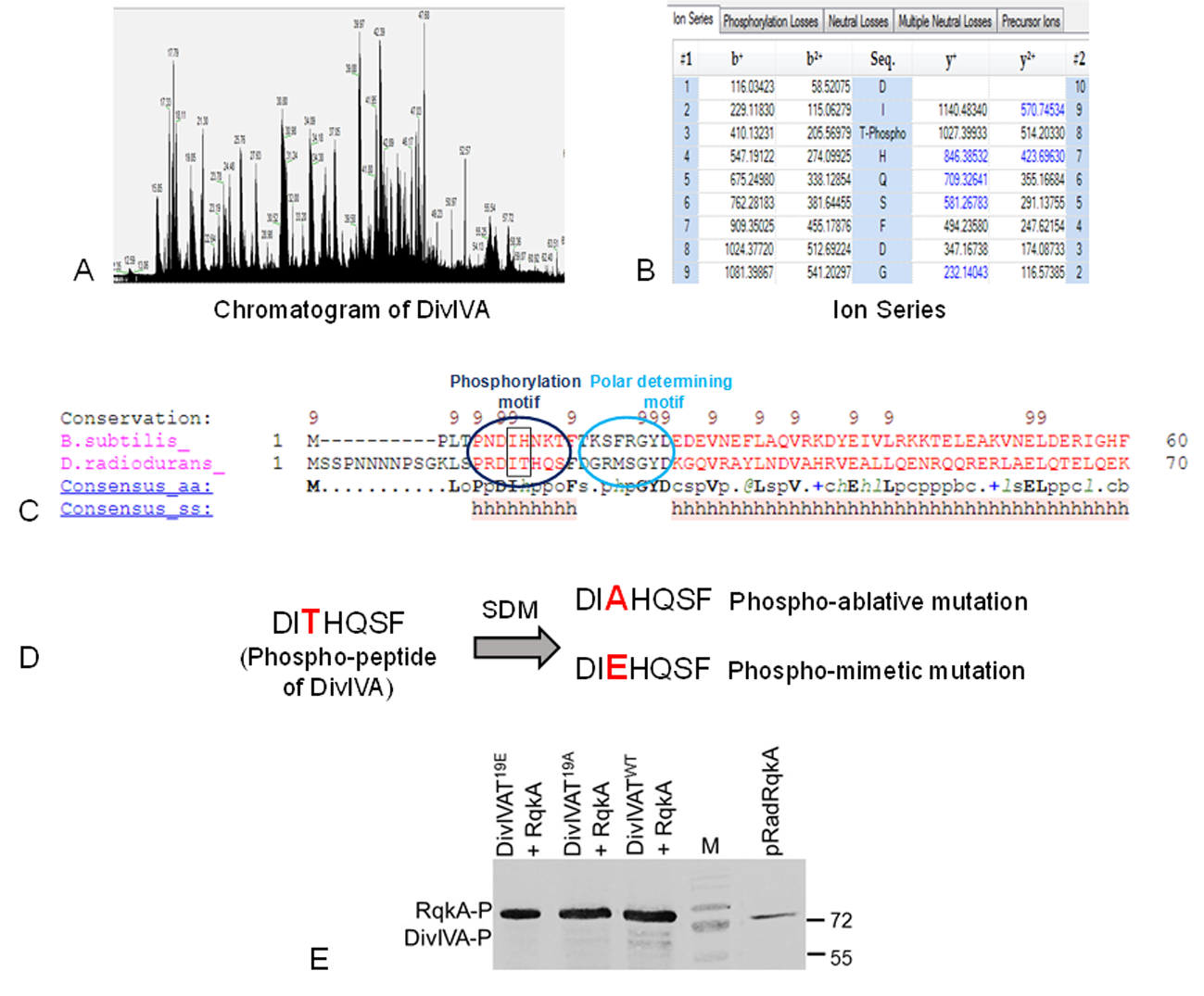

### Fig S2

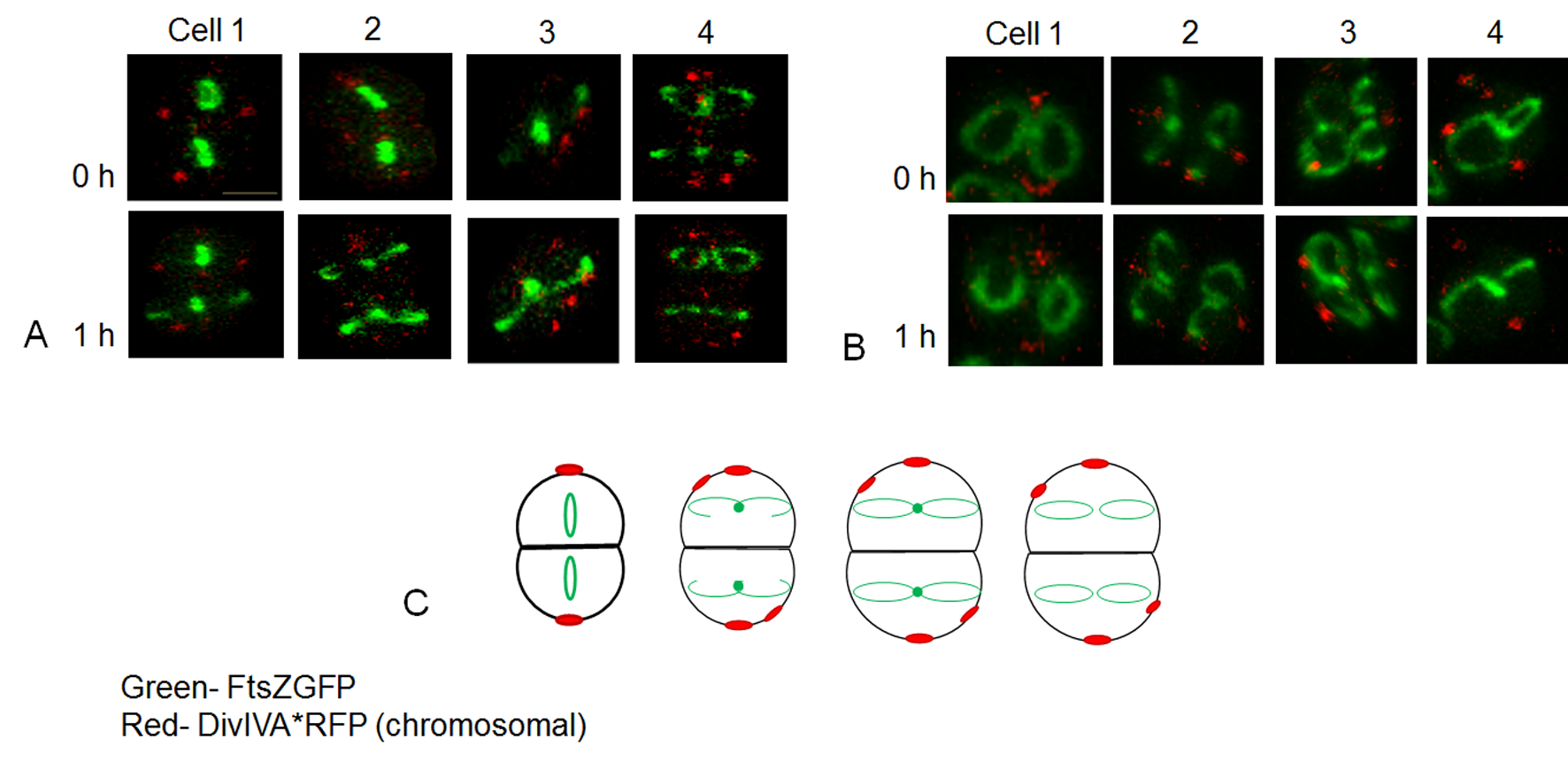

### Fig S3

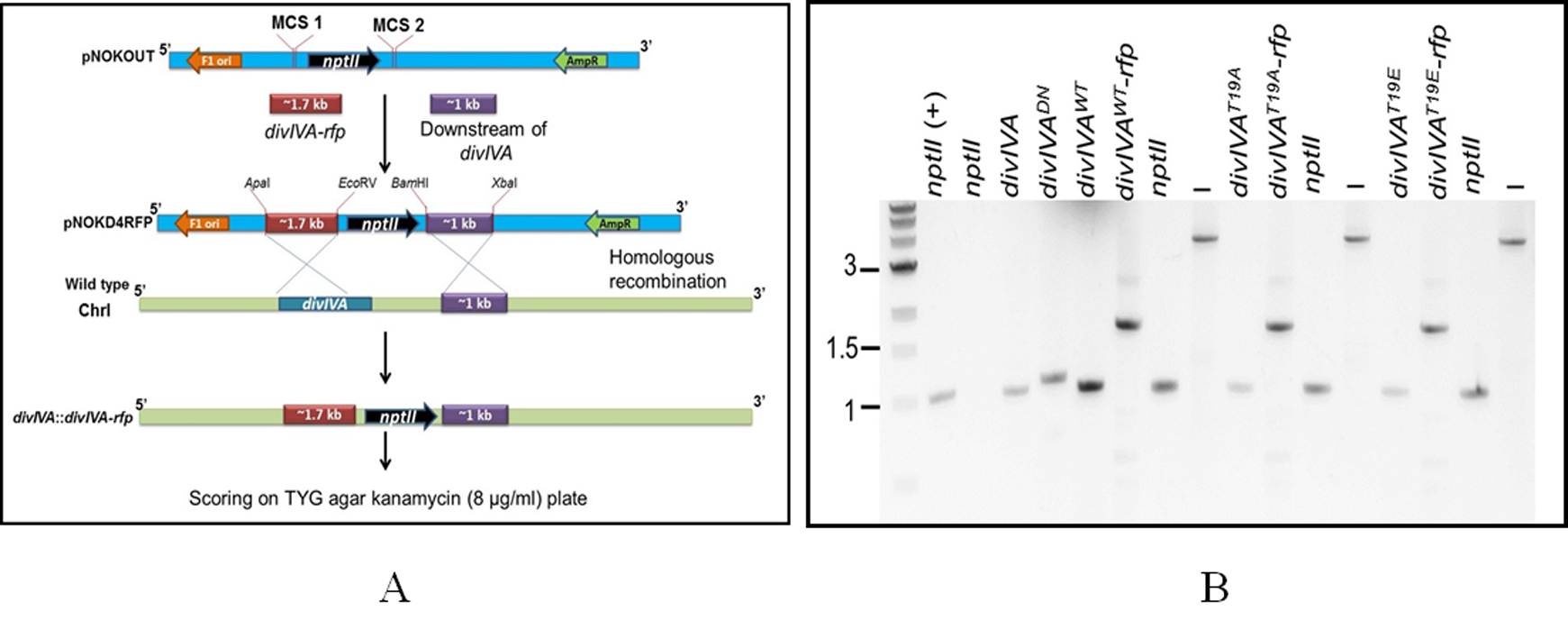
